# BioTrouble: A Multi-Agent Workflow for Troubleshooting Molecular Biology Techniques

**DOI:** 10.64898/2025.12.30.697016

**Authors:** Mehrdad Ameri, Hannie Yousefabadi, Amin Ramezani

## Abstract

Troubleshooting is a critical yet often underdocumented aspect of molecular biology experiments across laboratories. Failures in core techniques such as PCR, qPCR, molecular cloning, and related assays can lead to experimental failure, wasted resources, and delays in research progress. Here, we present BioTrouble, a multi-agent AI workflow designed to assist researchers in troubleshooting a wide range of molecular biology experiments. It leverages a custom-designed troubleshooting knowledge base through a retrieval-augmented generation (RAG) framework. BioTrouble employs small language models to generate the troubleshooting plan and utilizes a smart model routing system to manage cost per request. User interactions and feedback are stored as structured cases, enabling BioTrouble to expand its troubleshooting knowledge base and improve response generation over time. Compared with single-model SOTA LLM, BioTrouble generated comparable troubleshooting recommendations using small language models.

## Introduction

Troubleshooting is a critical component of molecular biology experiments. Experimental procedures frequently encounter unexpected challenges, requiring substantial time and effort to resolve. Failures in techniques such as PCR, qPCR, molecular cloning, and related assays are often addressed ad hoc, relying on the tacit knowledge and experience of individual researchers. It is worth to mention that not all researchers have the ability to perform troubleshooting (1–4). Numerous factors can contribute to cause issues in a single molecular laboratory test, and research and clinical laboratories routinely perform dozens of such assays. Identifying the root cause of every issue is a time consuming process, leading researchers to spend weeks or months seeking solutions. Rapid identification of solutions can yield significant savings in time, consumables, and researcher effort specially in life science projects. Artificial intelligence (AI), particularly AI agents, can be highly effective in this context by significantly accelerating troubleshooting (5).

AI agents are goal-oriented systems built on top of language models. They are capable of planning, acting, and interacting with tools or data sources to accomplish multi-step tasks, in contrast to conventional chatbots that only answer questions. The behavior of AI agents is governed by modular components, including system prompts, tools, memory, and routing logic, enabling precise customization for specific domains, such as molecular biology troubleshooting (6–9). Retrieval-augmented generation (RAG) represents a critical capability in these systems. Prior to generating a response, the agent can search for and retrieve relevant protocols, instrument manuals, or knowledge-base entries, and subsequently reason over this evidence rather than relying solely on internal parameters. This integration of tool use, domain-specific customization, and RAG, positions AI agents as superior to conventional chatbots for providing reliable, context-aware support in molecular biology workflows (10, 11).

Large language models function as the brain of AI agents (12). These models are available in various sizes and are generally classified as small or large according to their parameter count (13). A small language model is a term usually used for models below 10 billion parameters in size. While model scale and quality significantly affect an AI agent’s performance, employing the largest model is not always the most efficient approach (14). Assigning small language models to non-complex and routine subtasks, combined with effective context engineering, can achieve performance comparable to that of large language models (15). This approach enables the system to allocate large models to tasks that require advanced reasoning, while simpler operations are handled by smaller, more cost-effective models. As a result, this strategy reduces computational expenses and token consumption without compromising the overall reliability of the agent’s behavior (14). Various AI agents have been introduced for bioinformatics and biomedical applications including AutoBA (16), Bio-Copilot (17), BioRAGent (18), Biomni (19), DrugAgent (20), CRISPR-GPT(21), AutoProteinEngine (22). Here we introduce BioTrouble, a multi-agent workflow for troubleshooting molecular biology assays. It is a RAG-based AI agent workflow with the ability to take feedback from users and improve its response over time. The learning system implemented in BioTrouble is inspired by agentic context engineering (ACE) framework (23). BioTrouble has a model routing system that dynamically change the language model for each agent in the workflow due to the need to save token usage. It can work with a range of small language models to large language models considering simple to complex tasks. However, the primary objective of the BioTrouble architecture was to maximize the performance of small language models, including their ability to generate comprehensive troubleshooting plans. Evaluation results indicated that BioTrouble has a significant improvement in response compared to baseline models.

## Implementation

The overall framework of BioTroube is introduced in this section. Details about each agent, the retrieval system, the learning system are represented here. Additionally, the evaluation of BioTrouble against baseline models is outlined.

### Overall architecture of BioTrouble

BioTrouble is a multi-agent system built using the LangGraph framework. This state-based architecture integrates eight specialized agents that collaborate to process user requests, extract experiment information, retrieve relevant technical data, generate context-aware solutions, and control the workflow’s learning system. A central state management system coordinates data flow between these agents and maintains conversation history. To control token usage by each agent and computational efficiency, BioTrouble benefits from a model-routing system that handles dynamic model selection. A variety of small and large language models are assigned to each agent based on token usage and task complexity. Additionally, a learning system inspired by the ACE framework makes BioTrouble learn from user feedback during each conversation. Moreover, following some lab-specific troubleshooting guidelines, users can send their custom modifications and troubleshooting guides to BioTrouble, which will use that data to answer personalized questions in future conversations.

### Typical usage workflow

When a user starts conversation with BioTrouble, the router agent processes the user prompts and decides whether it is a chitchat, tip submission, or a technical laboratory question and controls the data flow in the downstream. The user can ask questions about protocol details, the troubleshooting guide, and other related questions to molecular biology experiments. If the initial question is ambiguous, BioTrouble asks for more information about the query. This helps the agent to answer better and more specifically. If the RAG needs to be activated during conversation, the retrieval agent obtains relevant documents from the database, and the agents downstream use those documents to generate a better response. For each solution and response generated by the agent, users can submit a helpful/harmful score with additional note. This feedback is stored and affects the agent’s response in the future for similar troubleshooting queries.

### Multi-agent implementation

BioTrouble is the orchestration of eight specialized agents:

- **Router Agent:** The router agent serves as the initial decision-making node for every user turn. Its primary function is to classify the user’s intent into main categories (work request, chitchat, tip submission) and sub-categories (e.g., troubleshooting, protocol, evidence, information). The agent employs a hybrid classification strategy to maximize reliability: it invokes a language model to semantically classify the statements, and if it fails, it attempts rule-based classification using regular expression (regex) patterns to detect high-confidence signals (e.g., specific keywords). The router also determines essential routing metadata, such as whether retrieval-augmented generation is necessary (e.g., for evidence requests or complex protocols) or if the conversation is a continuation of a previous thread, essentially controlling the downstream execution path.
- **Triage Agent:** This agent is responsible for structured context extraction. It functions as a semantic parser that converts unstructured natural language into a rigorous JSON schema used by downstream components. Using a language model, it extracts key experimental variables, including test type (e.g., PCR, Western Blot), specific symptoms (e.g., smear, no bands), and a detailed problem description. Crucially, it parses quantitative experimental conditions into a parameters object (e.g., temperature 95°C, concentration 2 mM), which allows direct validation against reference ranges. The agent maintains a cumulative state, merging new information with previously extracted context to handle multi-turn clarifications. It also generates clarifying questions if critical information is missing, which the response composer can later present to the user.
- **Retrieval Agent:** The retrieval agent orchestrates the acquisition of external knowledge. Unlike simple similarity search, it implements a two-stage “plan-and-execute” architecture. First, it uses a language model to generate an optimized retrieval plan, synthesizing a search query that targets specific symptoms and diagnostic patterns rather than generic terms. It also constructs flexible metadata filters (e.g., filtering by test type). Second, it executes this plan via the retrieval tool to retrieve related data from the knowledge base. The agent annotates the retrieved chunks, using a language model to summarize their relevance to the specific user problem, ensuring only high-quality evidence reaches the generator.
- **Generator Agent:** This agent serves as the brain of BioTrouble. It synthesizes the final troubleshooting plan or informational response by integrating the structured triage context, retrieved evidence, and session history. Additionally, it includes a “Failure Awareness” mechanism that checks the session history for actions that previously received negative feedback and explicitly instructs the model to propose fundamentally different approaches, preventing repetitive failure loops.
- **Response Composer Agent**: This agent is responsible for turning the system’s output into a clear and user-friendly answer. It takes the structured troubleshooting plan and makes it easy to read. The agent also helps users by summarizing earlier information and asking any follow-up questions needed for clarity.
- **Reflector Agent:** The Reflector Agent implements the “learning loop” of the system, inspired by the “Reflexion” and “ACE” (Agentic Context Engineering) frameworks. After an interaction, it analyzes the execution trace, user feedback (helpful/harmful signals), and failure reports. It uses a language model to generate a structured “reflection” that can identify the root cause of success or failure. The reflection outputs specific recommendations for knowledge base updates, such as modifying an existing bullet or creating a new one. This agent serves as a critic, converting raw interaction data into actionable insights to improve the system.
- **Curator Agent:** This agent organizes and standardizes proposed updates to the knowledge base. Acting as a bridge between the Reflector’s insights and the Delta Manager’s storage, it converts raw recommendations into Deltas (a request defining a specific change to the knowledge base (e.g., “change this rule”)). The Delta Manager acts as the execution layer and manages the lifecycle of these requests. It checks dependencies, resolves conflicts when multiple updates occur at the same time, and saves the final changes to the playbook.
- **Tip Ingestor Agent:** This agent captures user-submitted expertise and recommendations. It scans messages for explicit tips and uses a language model to translate free-text advice into the structured format required by the playbook (identifying test types, symptoms, and actions). Instead of modifying the database directly, it standardizes these contributions into formal update requests (Deltas).

### Retrieval-augmented generation (RAG) implementation

- **RAG knowledge base:** A JSON Lines file was created to store molecular biology lab techniques in bullet format. This knowledge base named playbook covers various molecular biology lab tests, categorize into the following sections: nucleic acid amplification tests (e.g., PCR, qPCR, RT-PCR), extraction and purification (e.g., DNA extraction, RNA extraction), sequencing-based methods (e.g. Sanger sequencing, NGS, pyrosequencing), hybridization-based assays (e.g., Southern blot, FISH, microarray), genotyping and mutation detection (i.e. RFLP, ARMS-PCR, fragment analysis), methylation and epigenetics (e.g., bisulfites conversion PCR, MeDIP-seq), RNA-based analysis (e.g., RNA-seq, single-cell RNA-seq), protein detection and analysis (e.g., Western blotting, immunoprecipitation), signal amplification-based analysis (bDNA), gel electrophoresis (agarose gel electrophoresis, and SDS-PAGE), molecular cloning and genetic modification (i.e. traditional cloning and CRISPR-Cas9 gene editing). Alternative names, the goal of the experiment, various applications, a detailed protocol, available commercial kits, and a troubleshooting guide have been provided for each experiment.
- **Strategic Query Synthesis:** Retrieval is not triggered by raw user input. Instead, the Retrieval Agent first analyzes the conversation state (symptoms, test type, and hypothesis) to generate a structured retrieval plan. This process synthesizes an optimized search query that strips away conversational noise and targets specific diagnostic markers. This step ensures that the search mechanism focuses on the underlying scientific phenomenon rather than the user’s phrasing.
- **Sparse Vector Modeling for Precision:** For knowledge representation, the system implements a custom in-memory vector store utilizing Term Frequency-Inverse Document Frequency (TF-IDF) rather than dense semantic embeddings. This design choice was driven by the need for high lexical precision in molecular biology, where specific alphanumeric codes carry exact meanings that dense models can sometimes conflate or “hallucinate” associations for. The system calculates Cosine Similarity between the query vector and document vectors to identify candidate evidence.
- **Multi-Factor Evidence Ranking:** Retrieved candidates undergo a ranking process to ensure the highest utility. The final relevance score for each knowledge unit (bullet) is calculated as a weighted function of three factors: i) Content Similarity: The TF-IDF cosine similarity score. ii) Metadata Alignment: A boost applied when the evidence matches the user’s specific context (e.g., matching test type or symptom tags). iii) Historical Utility: A dynamic “feedback score” derived from the learning system. Knowledge units referenced in successful troubleshooting sessions receive a weight boost, while those associated with negative feedback are penalized. This feedback loop ensures that the retrieval system effectively learns which playbook entries are most useful for specific problems over time.
- **Contextual Annotation:** Before data reaches the generation layer, a final filtration step is applied. The Retrieval Agent uses a language model to annotate the top retrieved chunks, explicitly marking their relevance to the current problem. This annotated context blocks the generic injection of non-pertinent facts, allowing the Generator Agent to reason strictly from high-quality, verified evidence.

### The learning system of BioTrouble

BioTrouble benefits from a learning mechanism inspired by the Agentic Context Engineering (ACE) framework. This system enables the agent to evolve its knowledge base over time, identifying gaps in its reasoning and correcting invalid troubleshooting protocols. The implementation is divided into three distinct architectural phases: reflective interaction analysis, atomic knowledge evolution, and consistency management.

- **Reflective Interaction Analysis:** The learning process begins with the Reflector Agent. Upon the conclusion of a troubleshooting session, it analyzes the interaction context. It uses a language model to generate a structured “Reflection” that infers the root cause of success or failure. By generating a structured Reflection, the system isolates the specific root cause of the error, converting raw conversation data into an actionable learning signal.
- **Atomic Knowledge Evolution (The Delta Protocol):** Knowledge updates are managed as transactional objects called “Deltas”. The Curator Agent translates the Reflector’s insights into these standardized payloads. Each Delta represents a discrete proposal (such as creating a new troubleshooting rule or modifying a relevance score) and carries necessary provenance data (source session ID, plan hash). The system is designed to automatically ingest non-conflicting updates. This allows the agent to patch its own knowledge base in near real-time, rapidly adapting to new user feedback.
- **Consistency Management:** The Delta Manager serves as the gatekeeper for these updates. It implements Identifier-Based Conflict Resolution. Before applying a Delta, it checks strictly for collision with other pending or recently merged updates targeting the same knowledge unit (bullet id). If multiple agents attempt to modify the same rule simultaneously, the manager detects this ID collision and moves the Delta to a “blocked” state for manual review. However, if no ID conflict exists, the Manager proceeds to the Auto-Merge pipeline, directly committing the change to the live playbook. This design prioritizes speed and responsiveness while preventing race conditions on specific database entries.

### Evaluation of BioTrouble

To evaluate the comparative performance of BioTrouble against baseline models, we employed the LLM-as-a-judge evaluation paradigm, a methodology that leverages the natural language understanding capabilities of large language models to assess qualitative dimensions of generated text (24, 25). Recent studies demonstrate that LLM-based evaluators can achieve high agreement with human expert assessments, with GPT-4 as a judge reaching 85% agreement with human experts, exceeding the human-human agreement baseline of 81% (24). To ensure evaluation fairness, we employed GPT-5.1, deepseek-v3.2, and gemini-3-flash-preview as three evaluators. All judge evaluations were performed using deterministic decoding settings (temperature = 0) to maximize reproducibility; however, complete determinism cannot be guaranteed due to the nondeterministic nature of hosted LLM inference. Three judges scored each query using set metrics, and the median score for each metric was calculated.

To ensure robust evaluation, we constructed a stratified test dataset comprising 200 queries categorized by difficulty (Easy, Medium, Hard). The dataset encompasses a diverse range of laboratory scenarios, spanning from direct informational lookups to complex, multi-step troubleshooting cases requiring deductive reasoning. Each question was paired with an expert-verified ground-truth answer to give judges a reference for scoring. The evaluation pipeline executed two parallel workflows for each query in the test set:

- The Challenger (BioTrouble): The agentic workflow was invoked (the learning system was disabled during this phase of the evaluation to maintain fairness in the assessment process).
- The Control (Baseline): The query was passed to a standalone baseline mirror of the generator agent model without access to external tools or the structured playbook (because the generator agent is the brain of BioTrouble, which designs the troubleshooting plan).

Both outputs were blindly evaluated by LLM Judges against a set of ground-truth points. The judges performed the evaluation based on the following metrics:

a. Subjective scoring metrics Across judge models *m* ∈ {1, …, *M*}:

- *a*_*m*_: judge *m*’s Accuracy score
- *C*_*m*_: judge *m*’s Completeness score

1. Accuracy (0 - 100): it measures the scientific validity of the response.

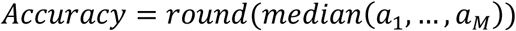
2. Completeness (0 - 100): it measures how well the response covers the ground-truth checklist and how practically useful it is for troubleshooting.

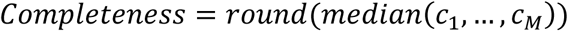
3. Subjective Quality (0 - 100): a composite metric defined as the arithmetic mean of Accuracy and Completeness. This primary Key Performance Indicator (KPI) serves as a holistic measure of troubleshooting effectiveness. 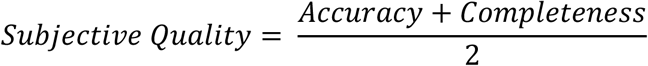 b. Objective scoring metrics:

- *G*: set of ground-truth points (e.g., {*GT*_1_, …, *GT*_*n*_})
- *C*: set of covered ground-truth points (*C* ⊆ *G*)
- *P*: set of predicted items extracted from the AI response (distinct items)
- *W*: set of incorrect keys (items in *P* judged wrong/unsafe/hallucinated)
4. Coverage (0 - 100): the percent of ground-truth (GT) points that the response covers. A GT point counts only if the judge includes its GT ID and provides a non-empty quote/snippet from the response as evidence.

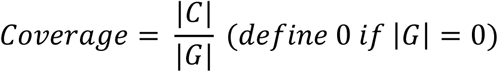

to represent it in percentage:

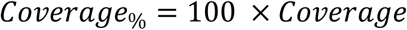
5. Wrong Rate (0 - 100): the percentage of predicted items in the response that are identified as incorrect, unsafe, or hallucinated.

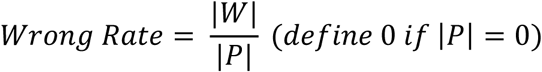

to represent it in percentage:

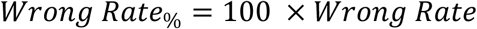
6. Purity (0 - 100): it measures how clean the successful coverage is among covered GT items and outright wrong items.

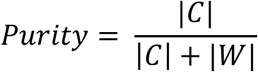

to represent it in percentage:

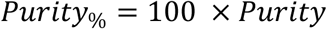
7. Integrity (0 - 100): it balances how much of the checklist is covered (Coverage) with how clean that coverage is (Purity). It is the harmonic mean of Coverage and Purity, so it is high only when both are good.

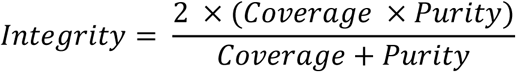

to represent it in percentage:

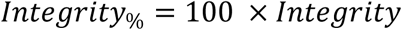
8. Objective Quality (0 - 100): it measures overall response quality using only auditable signals:

- how much of the ground truth troubleshooting checklist is correctly covered and,
- how rarely the agent makes wrong/hallucinated claims.

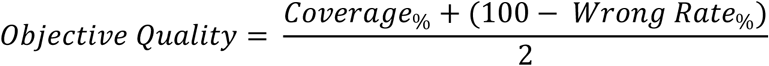

We evaluated four models and calculated all relevant metrics. BioTrouble aims to use SLMs for troubleshooting molecular biology experiments. To assess performance, we assigned three SLMs (Qwen3 8B, Llama 3.1 8B Instruct, and Gemma 2 9B) and one state-of-the-art LLM (GPT-4o) to the generator agent (which serves as the brain of BioTrouble) in separate runs. Each model’s performance in the multi-agent workflow was compared to its baseline performance outside the multi-agent setup. Given that BioTrouble uses a model-routing system, during the evaluation phase, we assigned only one model to the generator agent to maintain fairness in comparisons.

### Evaluation of learning system

To evaluate the BioTrouble learning system, we selected the top 20 most challenging questions from the benchmark dataset that the agent struggled with. Each question was given to the agent in 5 separate conversations. In each run, the agent received feedback with human supervision to activate the learning system. The generated answers from each run were qualitatively scored by a human expert on a scale of 1 to 10. The criteria for scoring the agent’s answer were the relevance of the answers to the question, the quality of details provided in each answer, and the likelihood of fixing the root cause. Finally, all scores were averaged across runs and converted to percentages.

### Cost per request assessment

One of the most important considerations in justifying the use of BioTrouble is cost. We tracked BioTrouble’s costs using the OpenRouter over 100 runs during the learning system evaluation, with all agents and learning components activated. These questions were also given to the GPT-4o model separately to calculate the average cost per request in BioTrouble, compared to a state-of-the-art large language model.

### User Interface

To facilitate user access to the multi-agent workflow, a Streamlit-based graphical user interface (GUI) was developed. This interface stores past conversations, displays active agents in real time, and allows users to send tips and feedback to agents via a chatbot.

## Results

### A multi-agent workflow for troubleshooting molecular biology experiments

Troubleshooting remains a central, time-consuming aspect of molecular biology workflows. To improve troubleshooting efficiency, we introduce BioTrouble, a multi-agent AI workflow. Figure 1 presents the general architecture of BioTrouble. Upon receiving a query, BioTrouble coordinates a sequence of actions among specialized AI agents to generate a comprehensive troubleshooting plan for the user. User messages can follow three paths in BioTrouble. The first path is chitchat, which controls messages that are not about molecular biology experiments or troubleshooting. The second path concerns work requests and is activated when the user asks about molecular biology protocols or troubleshooting. The third path is related to tip submission, which processes user recommendations about the experiment. These three paths are controlled by the router agent. BioTrouble uses a shared memory (conversation state), and all agents read from and write to a state object. Data is not passed directly from Agent A to Agent B; Agent A writes to the state, and Agent B reads from the state.

**Figure 1.**
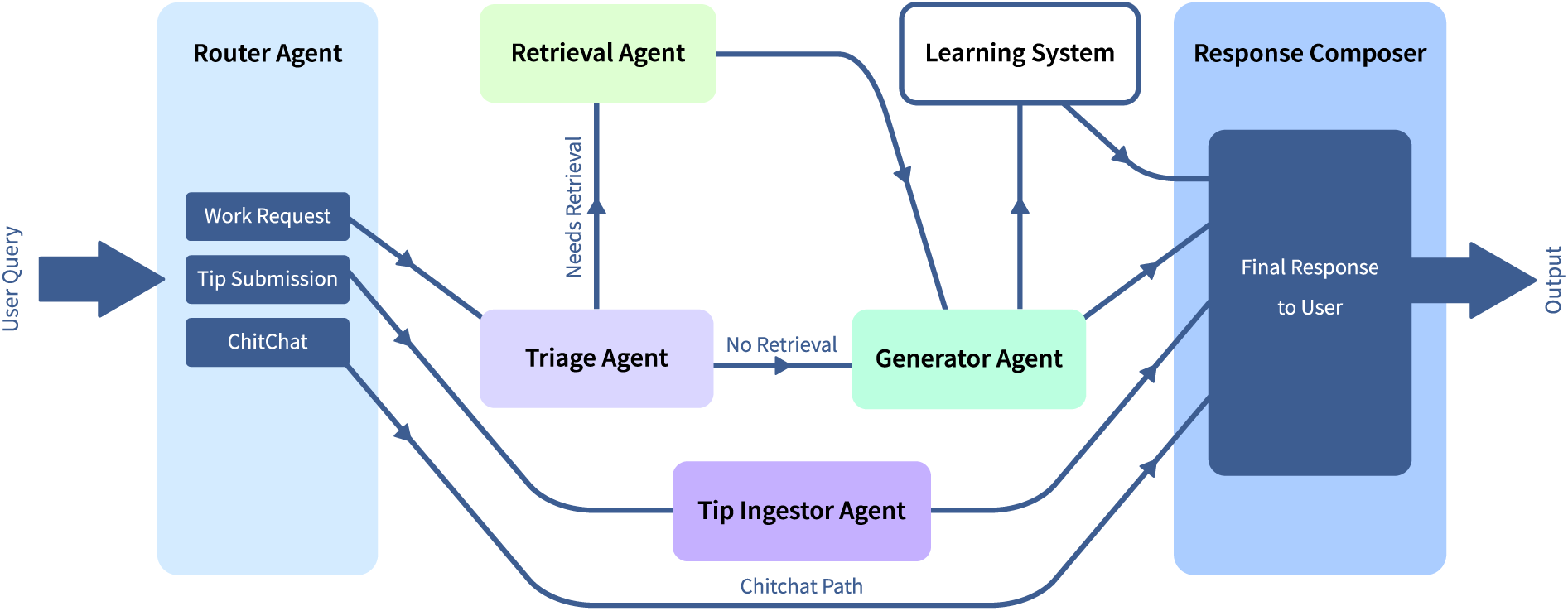
The general architecture of BioTrouble multi-agent workflow. The lines between agents are for simplifying data flow, and agents have no direct interactions with each other.

### All agents in BioTrouble have access to a shared state

BioTrouble employs a shared state object (Conversation State) as the central memory structure for inter-agent communication. Rather than passing data directly between agents, each agent reads from and writes to this common state, enabling a decoupled modular architecture. The state is empty at the start of each user session and progressively evolves as the workflow executes. Figure 2 represents the schematic overview of the state object. Upon receiving a user message, the Router Agent populates routing fields (e.g., router_decision). Subsequently, the Triage Agent adds structured diagnostic information (e.g., triage_result), and the Retrieval Agent appends ranked evidence (e.g., retrieval_results). The Generator Agent then synthesizes these inputs into a troubleshooting plan, and the state continues to evolve through downstream actions. Throughout this process, workflow control fields (e.g., visited_agents, loop_count, state_tokens) are continuously updated by the central coordinator to manage agent sequencing, prevent infinite loops, and inform dynamic model selection. This architecture ensures that every agent operates with full visibility into prior reasoning steps while maintaining a clear audit trail of the system’s decision-making process.

**Figure 2.**
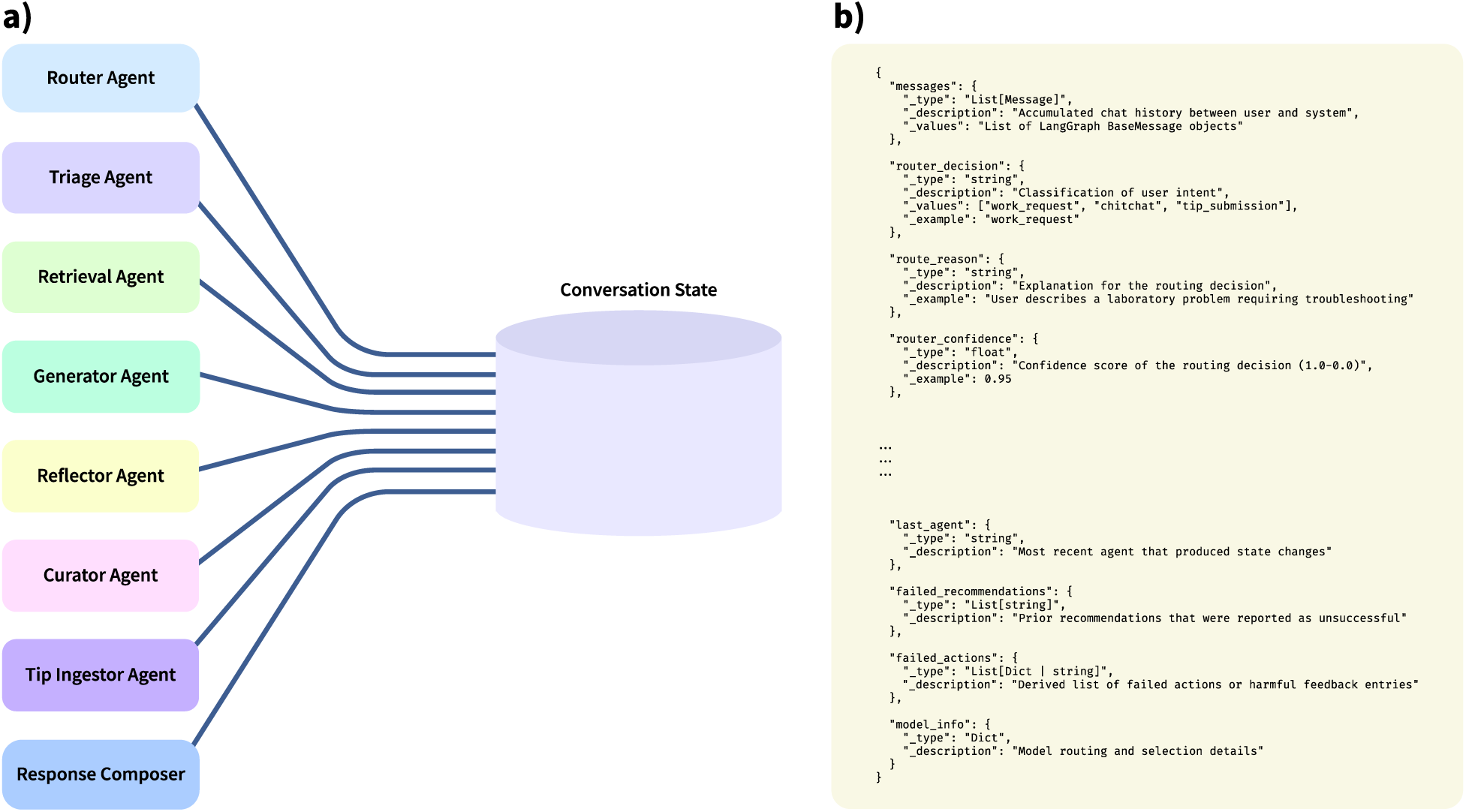
The state object of BioTrouble multi-agent workflow that evolves through workflow execution. a) graphical representation of how each agent is connected to conversation state, b) a JSON representation of state object (only part of state object is shown).

### A model manager assigns SLMs and LLMs to agents dynamically

To optimize the trade-off between cost and reasoning quality, BioTrouble benefits from a dynamic model routing system that assigns specific small or large language models to each agent based on the informational density of the conversation state. The system uses the total state token count as a quantitative proxy for task complexity. The architecture is defined by a hierarchical configuration in which each agent is provisioned with a tiered list of candidate models, triggered by specific token thresholds:

- Low-Density Task: For initial interactions, agents default to highly efficient SLMs. For instance, the Router Agent employs Granite-4.0-H-Micro for rapid intent classification of short utterances.
- High-Density Task: As the conversation deepens and the context window grows, the system automatically promotes agents to more capable lightweight LLMs (or LLMs due to

user needs). The Generator Agent, responsible for complex troubleshooting plans, switches from Llama 3.1 8B to GPT-4.1-mini once the context exceeds specified tokens, ensuring that complex reasoning is handled by the most capable model available when the information load is high.

The model configuration and token threshold we used for our setup were determined through a trial-and-error process, testing the quality of different models and thresholds. Users can update the model collections and token thresholds in the dedicated config file for this purpose as newer models are released. Table 1 lists the model options we used for each agent.

**Table 1.** The model collections for each agent in BioTrouble. For the generator agent, we have evaluated 3 small language models that produced reliable results. We call GPT-4o-mini SLM due to OpenAI’s official documentation. The parameter count of GPT-4o-mini is not officially declared, but it is called “Fast, affordable small model for focused tasks” in the model documentation available at https://platform.openai.com/docs/models/gpt-4o-mini. Additionally, we refer to GPT-4.1-mini as a Lightweight LLM because its parameter count is not officially declared, and it is not explicitly described as small.

| Agent | Model type | Model ID | Activation condition |
| --- | --- | --- | --- |
| **Router** | SLM | granite-4.0-h-micro | Low-Density Task |
|  | SLM | gpt-4o-mini | High-Density Task |
| **Triage** | SLM | qwen3-8b | Low-Density Task |
|  | SLM | gpt-4o-mini | High-Density Task |
| **Retrieval** | SLM | gpt-4o-mini | All Contexts |
| **Generator** | SLM | qwen3-8b,<br>llama-3.1-8b,<br>gemma-2-9b | Low-Density Task |
|  | Lightweight LLM | gpt-4.1-mini | High-Density Task |
| **Composer** | SLM | gpt-4o-mini | Low-Density Task |
|  | Lightweight LLM | gpt-4.1-mini | High-Density Task |
| **Curator** | SLM | gpt-4o-mini | All Contexts |
| **Reflector** | SLM | gpt-4o-mini | All Contexts |
| **Tip Ingestor** | SLM | gpt-4o-mini | All Contexts |

### BioTrouble benefits from a custom knowledge base to avoid hallucination

Retrieval-augmented generation (RAG) is a technique in which the LLM retrieves relevant evidence from an external knowledge base before generating its response, rather than relying only on its internal parameters. This can reduce the model’s hallucinations and produce a more reliable response. BioTrouble is a RAG-based agent with a custom-designed knowledge base that includes common molecular biology protocols and troubleshooting guidelines. The experiment categories and names in the BioTrouble knowledge base are listed in Table 2. This knowledge base is organized in a JSON lines file named playbook, and the retrieval agent can retrieve data from it. In addition to this file, user recommendations are organized into another JSON lines file similar to the main playbook, and the retrieval system retrieves data from both files.

**Table 2.** Molecular biology experiments included in RAG knowledge-base.

| Experiment category | Test included |
| --- | --- |
| **Nucleic Acid Amplification Tests (NAATs)** | PCR, qPCR, Reverse transcription PCR, Multiplex PCR, LAMP PCR, TMA PCR, RCA, NASBA |
| **Extraction And Purification** | DNA Extraction, RNA Extraction, Protein Extraction |
| **Sequencing Based Methods** | Sanger Sequencing, Next-generation DNA sequencing (NGS), Pyrosequencing |
| **Hybridization Based Assays** | FISH, Southern Blot, Northern Blot, Microarray |
| **Genotyping and Mutation Detection** | ARMS-PCR, RFLP, ASO probes, Fragment Analysis, MLPA, HRM |
| **Methylation and Epigenetics** | Bisulfite Conversion PCR, Methylation-Specific PCR, MeDIP-seq |
| **RNA-based Analysis** | miRNA profiling, RNA-seq, scRNA-seq |
| **Protein Detection and Analysis** | Western Blotting, Immunoprecipitation |
| **Signal Amplification-based Analysis** | Branched DNA Signal Amplification |
| **Gel Electrophoresis** | Agarose Gel Electrophoresis, SDS-PAGE |
| **Cloning and Genetic Modification** | Molecular Cloning, CRISPR-Cas9 |

### BioTrouble can learn and improve its knowledge base through user feedback and recommendations

ACE (Agentic Context Engineering) is an approach to improve an LLM’s performance by refining its context over time, rather than changing its model weights. It treats the context like a living playbook that keeps useful rules and avoids forgetting details. In BioTrouble, we wanted to improve agent response when RAG is activated. We designed the RAG knowledge base using a bullet-point structure inspired by ACE playbooks. In ACE, an evolving context is updated over time, whereas in BioTrouble, we decided to update the knowledge base based on user feedback, using a mechanism similar to the ACE framework. Therefore, BioTrouble has a feedback-driven knowledge refinement system. Rather than relying on static retrieval, the system continuously improves the relevance of its troubleshooting suggestions by dynamically adjusting the ranking of knowledge entries based on accumulated user feedback. When the agent provides a troubleshooting plan, the user evaluates its effectiveness. A “Helpful” vote serves as a positive signal, while a “Harmful” vote acts as a negative signal. These signals are recorded against the specific knowledge entry (bullet point) utilized in the plan. Every troubleshooting strategy carries persistent counters for helpful and harmful votes. This transforms the knowledge base from a static repository into a dynamic dataset where solution efficacy is explicitly tracked. The core improvement mechanism occurs during the retrieval phase of future conversations. When a new user query triggers a search:

- Prioritization: Strategies with a high net positive score (Helpful - Harmful) receive a significant boost in their retrieval ranking score. This ensures that proven solutions are presented first.
- Deprioritization: Strategies with negative feedback scores are penalized in the ranking, reducing their likelihood of being selected for future plans.

In addition to user feedback scoring, the system employs Reflector and Curator agents to expand the knowledge base. If harmful feedback is present, user-reported failure, or plan confidence < 0.6, the Reflector agent starts working and proposes edits such as adding notes or creating new bullets. The Curator then transforms these proposals into delta requests, which the Delta Manager integrates into the knowledge base. Over successive interactions, the system effectively learns by surfacing validated user solutions and filtering out ineffective ones, resulting in progressively higher-quality responses without model retraining. Figure 3 illustrates the learning system of BioTrouble.

**Figure 3.**
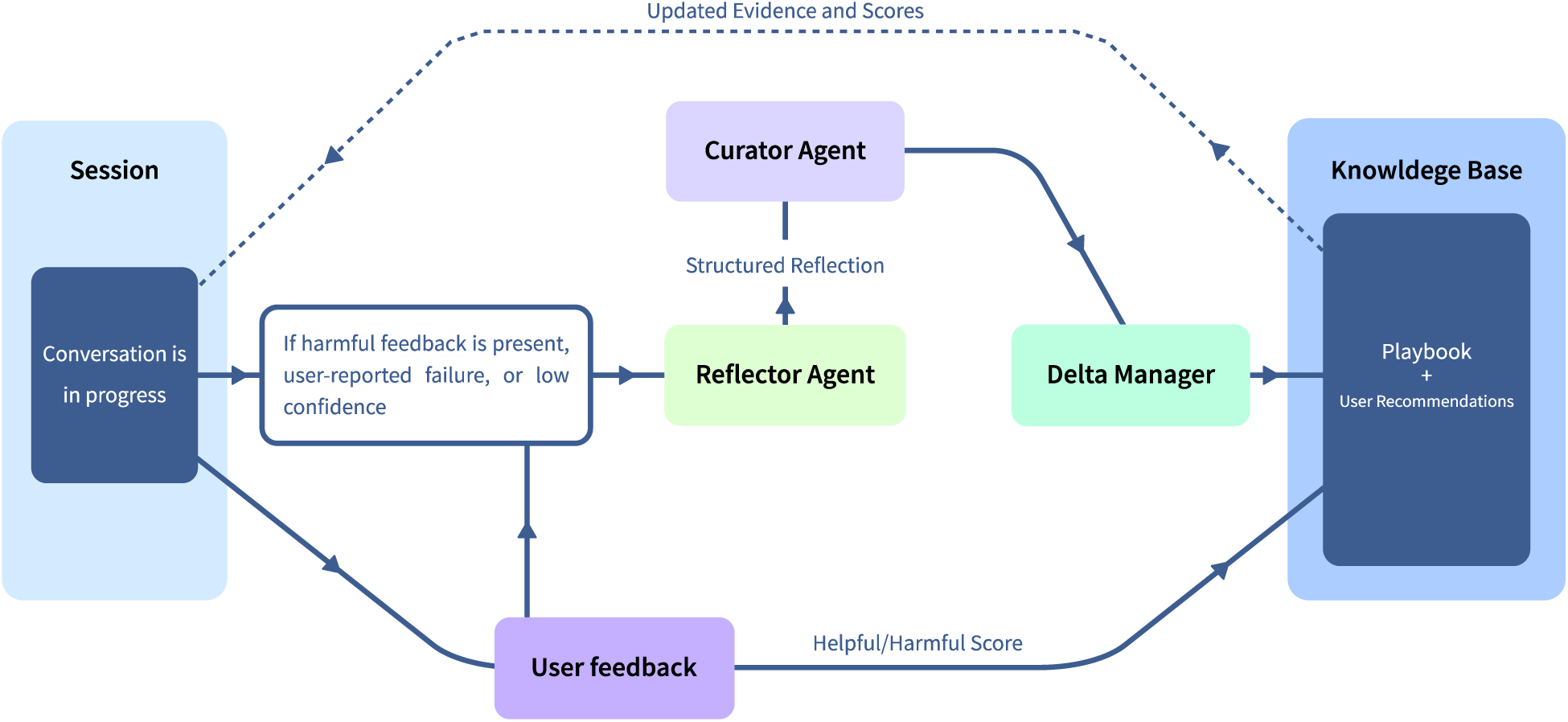
A representation of the learning system in BioTrouble.

### BioTrouble outperforms baseline model

In an LLM-as-a-Judge evaluation using 200 examples, BioTrouble beats the baseline model. In the BioTrouble multi-agent workflow, the generator agent serves as the core reasoning engine that generates a troubleshooting plan. We used a mirror of the model assigned to this generator agent as a baseline to compare its results with the full BioTrouble pipeline. Considering that one of the main goals of this project was to use small language models efficiently, we used Qwen3 8B, Llama 3.1 8B Instruct, 8-billion-parameter models, and Gemma 2 9B, a 9-billion-parameter model, for the Generator agent. In addition to SLMs, we have evaluated GPT-4o, a state-of-the-art LLM for this agent. The answer generated by these models in the multi-agent workflow achieved better overall scores than the baseline in all metrics we considered for this evaluation. Interestingly, in a categorical comparison (difficulty level of questions), the multi-agent achieved significantly better scores than the baseline across all four model evaluation runs, suggesting that BioTrouble can demonstrate its power under more challenging conditions. The comparison of the multi-agent workflow and the baseline model is shown in Figure 4.

**Figure 4.**
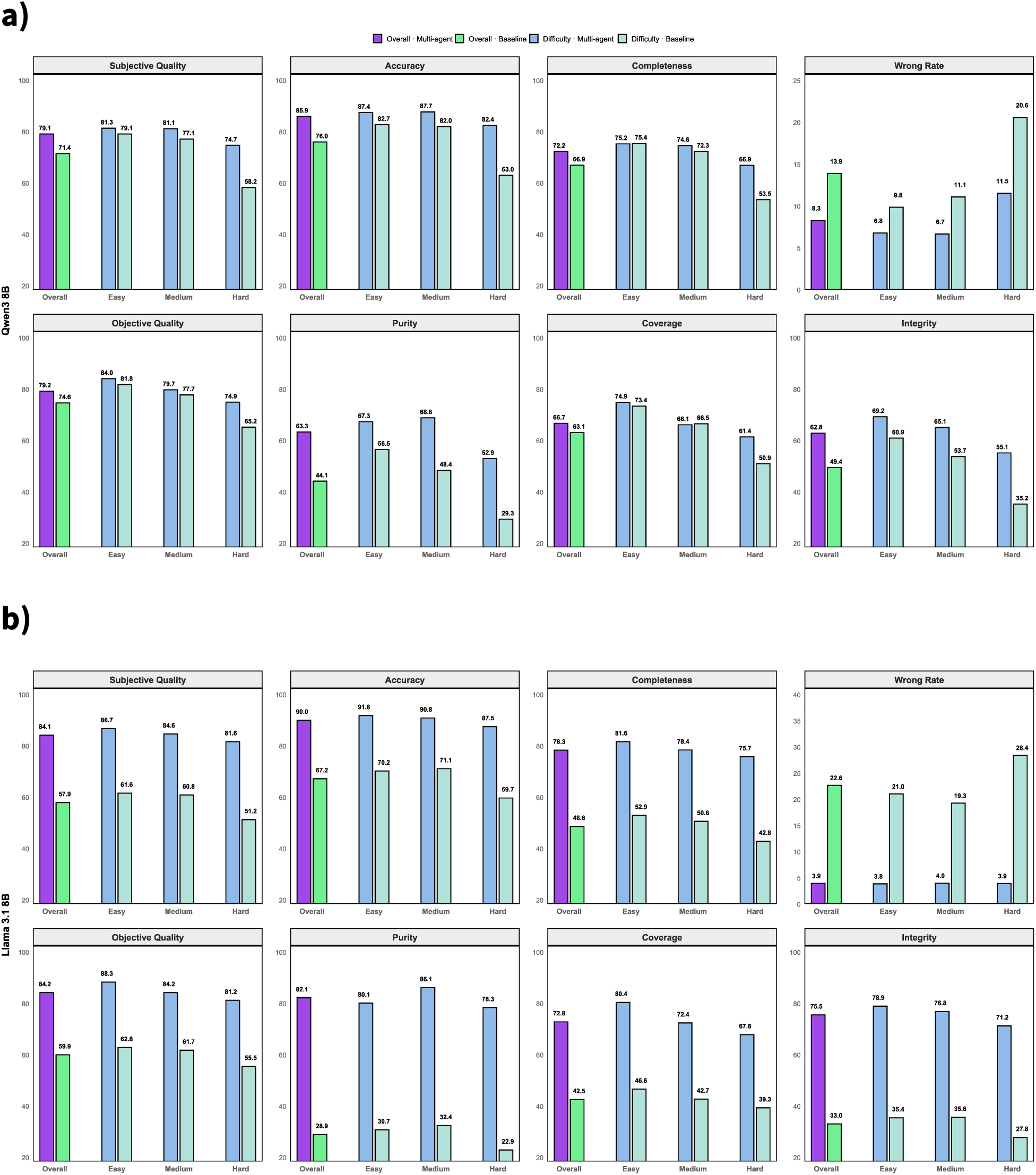
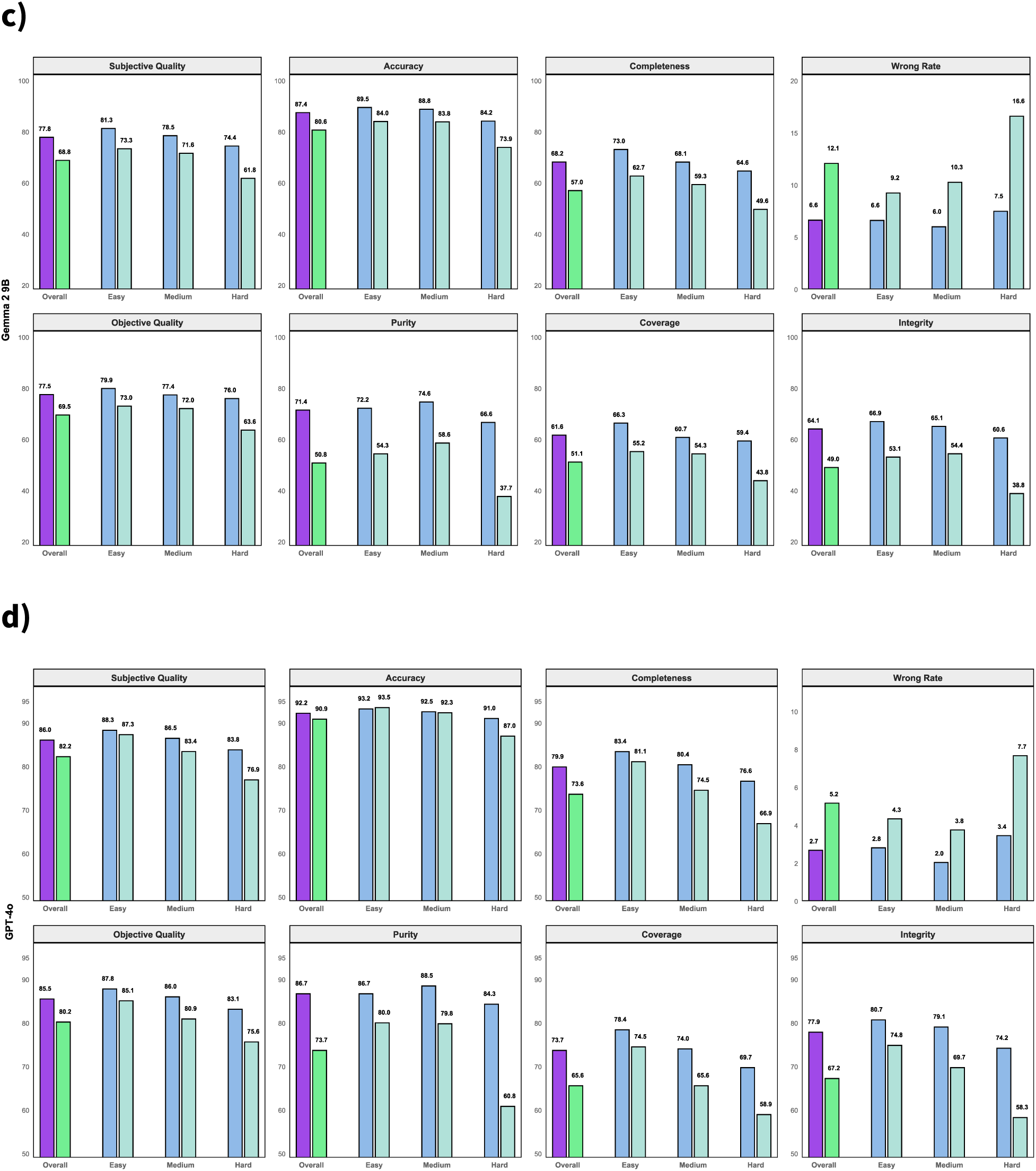
a) Qwen3 8B, b) Llama 3.1 8B, c) Gemma 2 9B, d) GPT-4o. For all metrics except Wrong Rate, higher values indicate better performance.

All three SLMs in the multi-agent setting achieved scores nearly good as the GPT-4o baseline. Interestingly, Llama 3.1 8B outperformed other SLMs across all metrics and beat the GPT-4o baseline in some of them (Figure 5).

**Figure 5.**
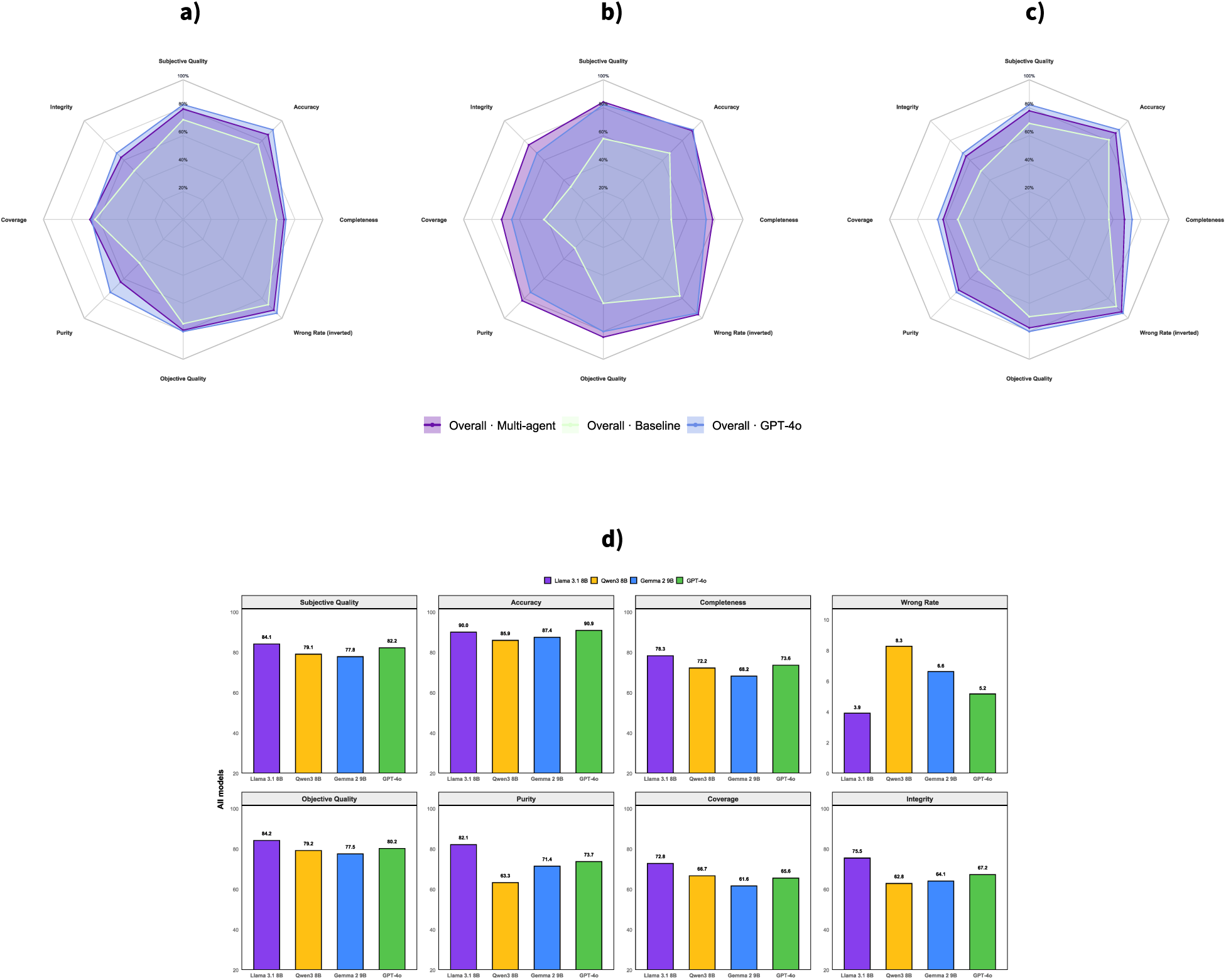
All three SLMs achieved comparable scores to the GPT-4o baseline. Because a lower Wrong Rate indicates better performance, unlike the other metrics, the Wrong Rate was inverted to improve visualization. a) Qwen3 8B, b) Llama 3.1 8B, c) Gemma 2 9B, d) All models.

### BioTrouble’s performance can be enhanced through the learning system

A key strength of BioTrouble is its learning system, which enhances the troubleshooting plan based on user feedback. To evaluate the BioTrouble learning system, the top 20 challenging questions that the agent struggled to answer during the main evaluation were selected. Each question was submitted to the agent in five separate conversations. With human supervision feedback, we observed improvement in BioTrouble’s response quality after five iterations. The evaluation of the learning system was performed by assigning Llama 3.1 8B to the generator agent. Figure 6 illustrates the evaluation of the learning system.

**Figure 6.**
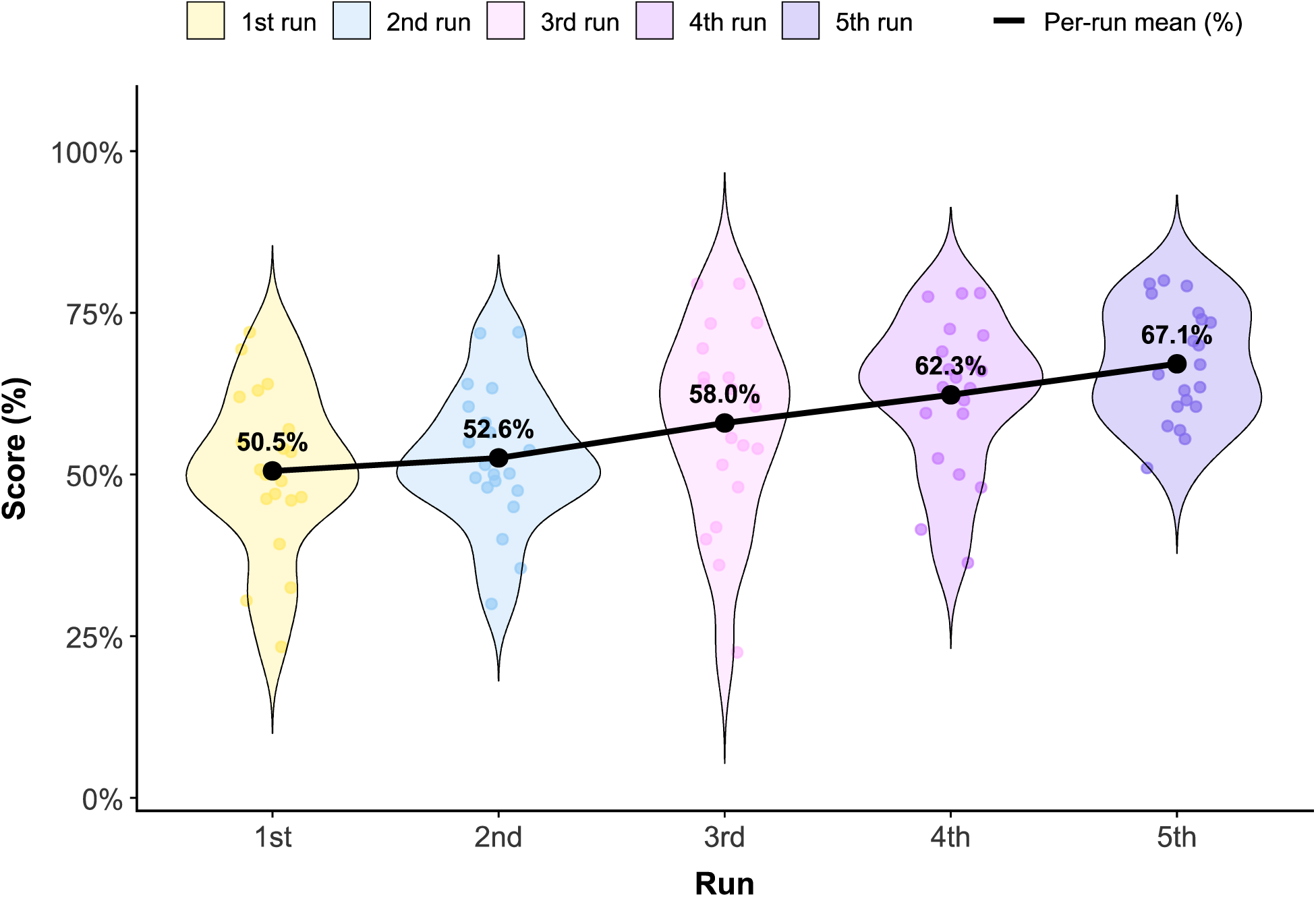
After 5 runs, the BioTrouble response improved through receiving user feedback.

### BioTrouble is more cost-effective than a state-of-the-art large language model (LLM) while offering comparable functionality

One reason to justify using BioTrouble is its cost per request. We recorded expenses for 100 requests using BioTrouble with all agents active via the OpenRouter API and repeated the process with the GPT-4o base model. BioTrouble averaged $0.0036 per request, while GPT-4o averaged $0.006. This 40% cost reduction, achieved while maintaining performance comparable to GPT-4o when Llama 3.1 8B was used as the generator agent.

## Discussion

Troubleshooting is a critical and often time-consuming part of molecular biology workflows. These experiments exhibit high sensitivity because they depend on precisely coordinated biochemical reactions and fragile biological materials. Minor deviations in material storage, reaction conditions, instrument calibration, or other assay-related factors can result in substantial impacts on specificity, yield, and reproducibility. Despite such issues being common, effective troubleshooting remains challenging to standardize across different laboratories and varying levels of expertise. To address this issue, we introduce BioTrouble, a multi-agent LLM-driven workflow for troubleshooting molecular biology experiments.

With the release of ChatGPT in 2022, large language models gained attention from the scientific community for use in specific tasks, including medical QA (26), clinical trials (27), clinical documentation and summarization (28, 29), and bioinformatics workflows (30, 31). Recently, AI agents have attracted researchers in the life sciences. Instead of relying solely on a language model, it is possible to achieve more reliable task execution in complex biomedical settings by providing a structured workflow that equips the model with planning, tool use, and memory. BioTrouble is capable of performing molecular biology experiment troubleshooting through the collaboration of multiple agents. Utilizing a collection of language models, BioTrouble achieved high scores across the metrics defined in this project.

Most AI agents for scientific applications use large language models as their first choice. As an example, in K-Dense, a multi-agent AI framework that automates scientific discovery for complex problems, including predicting biological age from transcriptomic data, they used Google’s Gemini 2.5 Pro as its core language model (32). In PrimeGen, a multi-agent AI framework designed to automate and assist primer design for targeted next-generation sequencing, researchers employed GPT-4o as the core language model to coordinate multiple specialized agents in this workflow (33). However, large language models are not the best option in most cases. Small language models with fewer than 10 billion parameters can perform specific tasks with reasonable efficiency when a good architecture is implemented (14). In BioTrouble, one of our main goals was to provide a system that leverages the power of SLMs to efficiently troubleshoot molecular biology experiments. Our results showed that Qwen3 8B, Llama 3.1 8B Instruct, and Gemma 2 9B, three SLMs assigned to the brain of BioTrouble, along with other agents controlling data flow, can outperform their baselines. Furthermore, they could achieve high performance comparable to a state-of-the-art LLM, GPT-4o. This shows that LLMs are not always the best solution, and we can achieve reasonable performance with SLMs when good context engineering is implemented. Furthermore, our evaluation indicates that BioTrouble, even with a larger model like GPT-4o for its brain (the generator agent), can outperform the baseline model.

One of the main issues in working with language models is their limited internal knowledge, which limits their ability to adapt to a specific task. Two popular approaches are used to overcome this issue. The first one is fine-tuning language models on a specific dataset, which is time-consuming and requires powerful infrastructure. Second is RAG, which provides an external knowledge base for the model to review before generating an answer. In BioTrouble, we implemented a RAG knowledge base with specific protocols and guidelines for troubleshooting across several molecular biology experiments. A major advantage of BioTrouble is its learning system, which allows users to improve the RAG knowledge base through submitting feedback in different conversations. Our results showed that, across five iterations in different conversations, BioTrouble can generate a better troubleshooting plan based on user feedback. This makes it a dynamic system that can provide customized answers in various laboratories. Additionally, because it can handle user tip recommendations and use them to answer future similar questions, BioTrouble can improve troubleshooting efficiency in projects and laboratories that are sensitive to their protocols.

A major concern in developing an AI agent is the cost of usage. Through our evaluation, we found that BioTrouble can provide a troubleshooting plan with 40% lower cost than GPT-4o and a performance nearly as good as this SOTA LLM. The model routing system in BioTrouble assigns different language models to agents based on token usage, thereby significantly affecting the cost per request. We used an OpenRouter API to run and monitor our evaluation, but users can also use open-source models locally and offline. This can result in significant cost reduction and ensure the safety of sensitive data.

A primary limitation of this project is the availability of free high-quality data. To build the knowledge base, we carefully gathered protocols and troubleshooting guidelines from sources licensed under Creative Commons Attribution (CC BY). Due to the quality of the knowledge base, the troubleshooting plan provided by BioTrouble can be considerably improved. Additionally, because our research goal was to build a system around small language models for troubleshooting, we didn’t cover all existing molecular biology experiments in the knowledge base. By expanding the BioTrouble knowledge base to include additional molecular biology tests, more researchers can use this project. Another limitation of this project is the small evaluation dataset. We used 200 questions for the evaluation, which represents a small fraction of the troubleshooting scenarios and questions related to the molecular biology experiment. We aim to expand the evaluation dataset in the future to examine BioTrouble’s capabilities with a more diverse set of questions.

## Conclusion

In conclusion, BioTrouble addresses a key challenge in molecular biology by offering a multi-agent workflow that accelerates experiment troubleshooting across laboratories and expertise levels. By integrating planning, tool use, memory, and a RAG knowledge base with a learning system, BioTrouble offers a practical path to more reliable, customizable troubleshooting over time. Our results demonstrate that strong troubleshooting performance does not require an expensive large language model; effective context engineering and agent collaboration allow small language models to perform competitively. While this study is limited by the scope of experiment coverage and evaluation size, these constraints identify clear opportunities for future expansion. Enhancing the knowledge base and extending evaluations to a broader range of assays and scenarios will further advance BioTrouble as a dynamic and generalizable platform for molecular biology troubleshooting.

## Author contributions

M.A. designed and implemented the architecture and code of BioTrouble, H.Y. designed and generated the knowledge base, and A.R. supervised the project. All authors contributed to the BioTrouble evaluation and to writing the project manuscript.

## Code and data availability

All code and data related to this project are available at GitHub: https://github.com/mehrdadameri/BioTrouble.

